# Evidence of Increased Hemangioblastic and Early Hematopoietic Potential in Chronic Myeloid Leukemia (CML)-derived Induced Pluripotent Stem Cells (iPSC)

**DOI:** 10.1101/2020.06.17.140202

**Authors:** G. Telliam, O. Féraud, S. Baykal-Köse, F. Griscelli, J. Imeri, T. Latsis, A. Bennaceur-Griscelli, A.G. Turhan

## Abstract

Hemangioblasts derived from mesodermal lineage are the earliest precursors of hematopoietic stem cells and endothelial cells. Embryonic stem cells (ESC) and induced pluripotent stem cells (iPSC) are the only experimental systems in which these cells can be assayed and quantified. We show here using CML-derived iPSC and blast-cell colony forming (Bl-CFC) assays that hemangioblasts are highly expanded in CML derived iPSC as compared to human H1-ESC-derived hemangioblasts. BCR-ABL signaling pathway is intact in these cells with evidence of CRK-L phosphorylation which is reduced by the use of Imatinib. Hematopoietic progenitor assays generated using blast-CFC demonstrates also a highly increased hematopoietic progenitor potential of these cells as compared to H1-ESC. The same results were also obtained using hematopoietic progenitor assays via embryoid body formation. In CML iPSC, we have also found a significant reduction of Aryl Hydrocarbon Receptor (AHR) expression which is involved in hematopoietic quiescence. Further inhibition of AHR using StemRegenin (SR1), an AHR antagonist, led to an increase of blast-cell colonies in CML iPSC whereas the use of an AHR agonist inhibited blast cell colonies. Thus, our results show for the first time, the possibility of establishment of a myeloproliferative phenotype using patient-derived iPSC and the presence of a major expansion hemangioblast compartment and derived hematopoietic progenitors in this context. They also suggest that the AHR signaling pathway could represent a novel druggable target in CML.

## INTRODUCTION

Chronic myeloid leukemia is the prototype of hematopoietic stem cell malignancy involving a highly primitive hematopoietic stem cell (HSC). The disease presents essentially by the expansion and proliferation in peripheral blood, of differentiated cells and hematopoietic progenitors of myeloid lineage but lymphoid B cells lineage can also be involved (Martin et al., 1982). There is a controversy with regard to the involvement of endothelial lineage in CML which would suggest that a stem cell with both hematopoietic and endothelial potential would be the target of transformation by BCR-ABL (Fang, 2005; Otten et al., 2008).

Patient-derived induced pluripotent stem cells (iPSC) represent a powerful tool to model several types of diseases (Zeltner and Studer, 2015) and especially chronic myeloid leukemia (Bedel et al., 2013; Hu et al., 2011; Kumano et al., 2012; Suknuntha et al., 2015). We report here for the first time that the earliest common hematopoietic and endothelial precursor designed as hemangioblast is highly expanded in stem cells assays using iPS cell lines derived from a patient with CML. We show that CML-derived iPSC generate high numbers hemangioblasts detected by blast-cell colony forming cell (bl-CFC) assays. Likewise hematopoietic progenitors derived from either bl-CFC or embryoid bodies are also highly increased in CML-derived iPSC. Finally we show the potential of manipulating this expansion by using AHR pathway.

## MATERIALS AND METHODS

### Patient-derived iPSC and characterization

Generation of CML iPSC has been previously described (Telliam et al., 2016). Briefly, peripheral blood cells were obtained at diagnosis from a 14-year old CML patient with informed consents according to the declaration of Helsinki. This patient was treated with Imatinib mesylate as a first line therapy. At +12 months post-therapy, there was no molecular response and no other TKI was available in 2004. An allogenic bone marrow transplant was performed. The patient remains in complete remission since the transplant performed in 2004. To generate iPSC, leukemic CD34+ isolated and cryopreserved at diagnosis were used as previously described (Telliam et al., 2016). All analyses were performed using a polyclonal stock of iPSC. This cell line designed as PB32 was characterized by using in vitro and in vivo pluripotency tests as previously described (Telliam et al., 2016).

### iPSC and ESC cultures

hIPS cell lines were maintained on the mitomycin-C-inactivated mouse embryonic fibroblast feeder cells with DMEMF12 supplemented with 0.1mg/ml bFGF and treated 2h with 1mg/ml Collagenase type IV (gibco by life technologies ref 17104-019). H1 cell line was used as previously described, after the authorization obtained from Agence de Biomedicine (Bennaceur-Griscelli, 2007). For cultures with Imatinib mesylate, cells were cultured for 5 days at 1μM before protein extraction.

### Embryoid body assays

For Embryoid Body (EB) formation, ES and iPS cells, at day 6–7 after cell passage, were treated with collagenase IV. Clumps were cultured in Iscove’s modified Dulbecco’s medium (IMDM, Invitrogen) supplemented with 1% penicillin/streptomycin, 1 mM L-glutamine, 15% fetal calf serum (FCS, Invitrogen), 450 μM monothioglycerol, 50 μg/mL ascorbic acid (Sigma Aldrich) and 200 μg/L transferrin (Sigma Aldrich) and supplemented with hematopoietic cytokines: 100 ng/mL stem cell factor (SCF), 100 ng/mL fms-like tyrosine kinase 3 ligand (Flt-3L) and 50 ng/mL thrombopoietin (TPO) (all from Peprotech). ES and IPS derived EB were cultured in ultra low attachment 6-well plates (Costar) for 16 days. Media was changed two or three times depending on EB proliferation. All cultures were incubated at 37°C in 5% CO2. For cultures with Imatinib mesylate, cells were cultured for 16 days at 1μM before protein and RNA extraction.

### Blast-colony forming assays

To obtain Bl-CFC, iPSC pellet was resuspended in Stemline II Hematopoietic Stem Cell Expansion medium (Sigma-Aldrich S0192) supplemented with 1% peni-streptomycin and L-glutamine, 50ng/ml rhVEGF (Peprotech) and 50ng/ml rhBMP4 (Peprotech). After 48h we added 20ng/ml of rhSCF (Peprotech), rhTPO (Miltenyi Biotec) and rhFLT3L (Peprotech). The embryoid bodies obtained at day 3.5 were collected, dissociated with pre-heated stable trypsin replacement enzyme TrypLE Express (Gibco by life technologies ref 12605-10) and filtered with 40μm Nylon Mesh sterile cell strainer (Fisher Scientific). Cells were suspended in Stemline II Hematopoietic Stem Cell Expansion medium (Sigma-Aldrich S0192) supplemented with 1% peni/streptomycin and L-glutamine, 50ng/ml rhVEGF, rhBMP4, rhFLT3L and rhTPO, 20ng/ml FGFb (Peprotech) and 5Units/ml EPO (Peprotech) and transferred in Methocult SF H4436 (Stem cell technologies) for 5 to 7 days.

### Hematopoietic differentiation

CFC assays from ES and iPS cells from EB or BL-CFC were performed by plating 10×10^3 cells/mL into MethoCult GF (H4435 StemCell Technologies) in triplicate. For ITK conditions, MethoCult GF were supplemented with Imatinib mesylate (1μM) or Dasatinib (5nM). Hematopoietic colonies (BFU-E, CFU-M, CFU-G, CFU-GM and CFU-Mixtes) were counted at day+14 according to standard morphological criteria. CFC assay was performed by the same manipulator during all clonogenic differentiation experiments.

### RNA and Q-RT PCR analyses

Total RNA was isolated using TRIzol (Ambion from Life technologies ref 15596026) extraction protocol and treated with DNAse I (Invitrogen). cDNA was synthesized using affinity script multi temperature cDNA synthesis kit. PCR amplification was conducted with Fast start DNA polymerase (Roche) or Platinium taq (Thermofisher).

For RT-PCR-mediated BCR-ABL detection, we used the European Leukemia Network primers (BCR-F: TGACCAACTCGTGTGTGAAACTC; ABLK-F: CGCAACAAGCCCACTGTCT; ABLK-R: TCCACTTCGTCTGAGATACTGGATT) The other primers used are listed in Table 2.

RTqPCR amplification was conducted with TaqMan Universal PCR Master Mix No AmpErase UNG (Applied Biosystems). Amplification products were run in a 1% or 0,8% agarose electrophoresis gel and detected with the GenoView UV Transilluminator. Unpaired t.test with Welch’s correction was performed from Prism software.

### Phenotypic analysis of hematopoietic progenitors

Hematopoietic cells were obtained using both via EB and Bl-CFC generation. To analyze hematopoietic cells generated via EB differentiation, cells were dissociated with pre-warmed stable trypsin replacement enzyme TrypLE Express (gibco by life technologies ref 12605-10) and filtered with 70μm Nylon Mesh sterile cell strainer (Fisher Scientific). Living cells were counted using trypan blue and stained.

To generate hematopoietic cells from Bl-CFC, we used colonies at day +5 of differentiation. Bl-CFC on methylcellulose were washed 3 times with PBS. The pellet was resuspended in Iscove’s MDM medium and filtered with 70μm Nylon Mesh sterile cell strainer (Fisher Scientific). Living cells were counted in trypan blue and stained with the following antibodies in PBS 4%BSA at 4°C 45 min: CD45Pe-Vio770 (Miltenyi), CD34Vioblue (Miltenyi), CD41aPE (BD), CD43FITC (Miltenyi), CD309APC (BD), CD31FITC (Beckman Coulter), CD235aAPC (BD), CD144 FITC (Miltenyi), BB9PE (BD). After 1h, cells were washed and resuspended in PBS 4%BSA with viability staining reagent 1μg/ml (7-AAD) 7-aminoactinomycin D (Sigma Aldrich). Stained cells were analyzed with a MACSQuant 10 (Miltenyi Biotec) flow cytometer and Flowjo analysis software.

### Western Blots

The cells were lysed in ice with RIPA buffer containing (NaCl [200mM], Tris [pH 8; 50mM], Nonidet P40 [1%], acide deoxycholate [0.5%], SDS [0.05%], EDTA [2mM]) supplemented with 100μM phenylmethylsulfonyl fluoride (PMSF), 1mM sodium fluoride (NaF), 1mM orthovanadate (Na_3_VO_4_). Separation of proteins was performed by electrophoretic migration on a 3-8% polyacrylamide gel under denaturing conditions. Proteins were then transferred in a semi-liquid condition on PVDF membrane pre-activated in methanol. After saturation with TBS Tween 5% BSA for 1h and hybridization of the membranes with primary overnight (BCR-ABL Ab3 Calbiochem, pCrkL Tyr207 Cell Signaling, pTYR 4G10 Millipore) and secondary 1h antibodies coupled to HRP, the signal was revealed by chemiluminescence with SuperSignal West Dura or Femto reagents and acquire with G:BOX iChemi Chemiluminescence Image Capture system.

### AHR experiments

They were performed with the goal of detecting the effect of SR1 addition during the generation of Bl-CFC and during the hematopoietic progenitor generation (Figure 9). Bl-CFC assays were started in iPSC in the presence or absence of SR1 1μM.

At day + 3.5 of Bl-CFC differentiation, cells were dissociated and divided in 3 separate conditions for CFC assays in MethoCult, including control, sample supplemented with 1μM SR1 (Sigma Aldrich) or seeded with 100nM FICZ (Sigma Aldrich). After 14 days the methylcellulose were washed and cells counted with Trypan blue.

### Statistical analyses

Unpaired t.test and Two way ANOVA statistical analysis were performed with PRISM software from at least 3 experiments.

## RESULTS

### BCR-ABL expression in CML iPSC

BCR-ABL expression was analyzed in CML IPSC at pluripotent stage by first with Western blots in the presence of appropriate controls which included iPSC line PB33 (iPSC without Ph1 chromosome), K562cells, the UT7 cell line and its counterpart expressing BCR-ABL (UT7/11) as well as hESC line H1. As can be seen in Figure 1A, BCR-ABL was not detected in control embryonic stem cell line H1, IPS cell line PB33 and hematopoietic cell line UT7 (**figure 1A**). The CML iPSC PB32 cells expressed BCR-ABL protein but at much lower levels as compared to CML cell lines K562 and UT7-11. The analysis performed after Imatinib Mesylate (IM) treatment did not reveal any difference in protein expression. A quantification of BCR-ABL gene expression levels with and without IM in CML-IPSC showed no significant difference (**figure 1B**).

**Figure 1:**
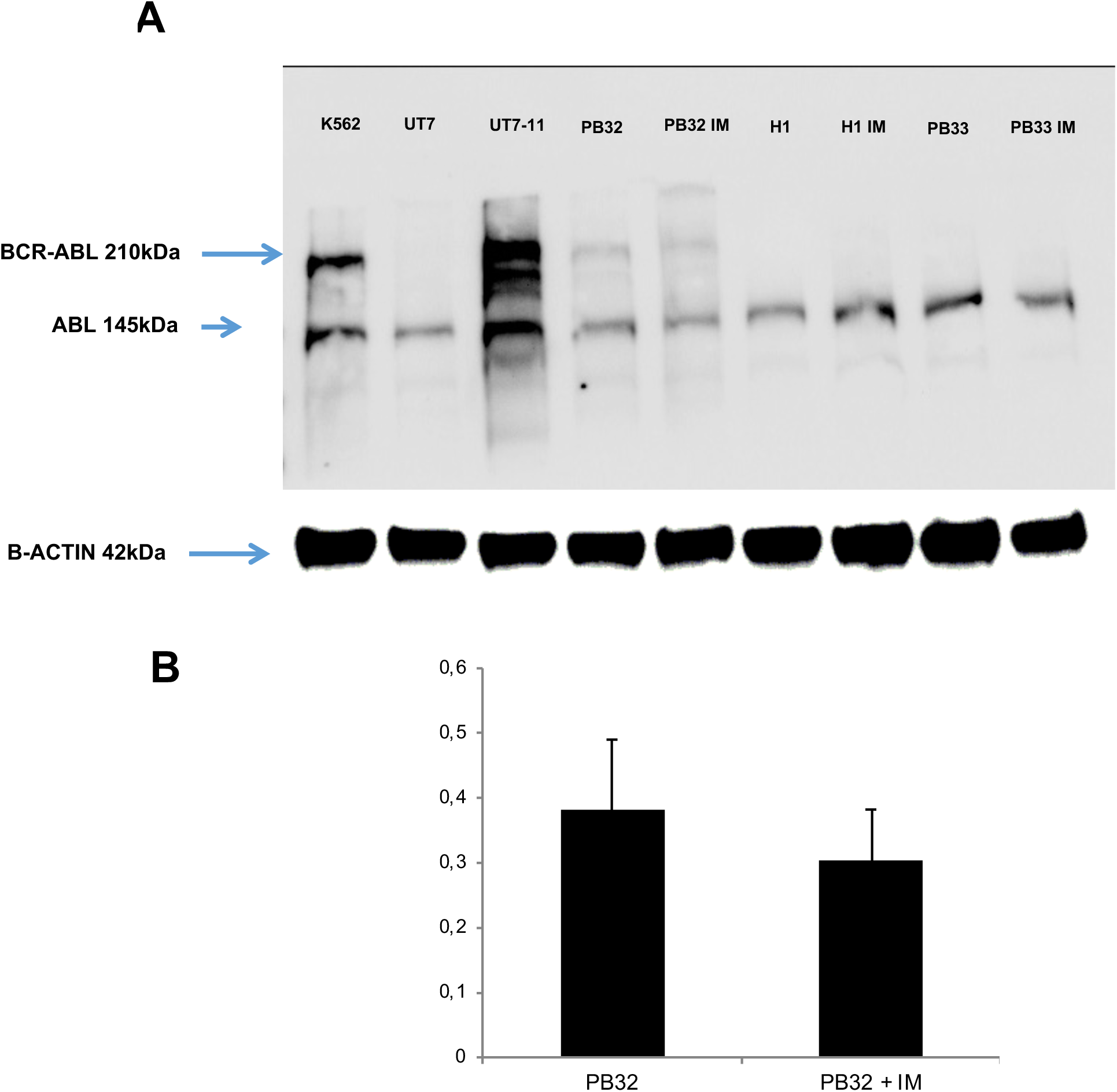
BCR-ABL expression in CML iPSC line PB32. **(A)** Western blot analysis for evaluation of BCR-ABL expression. in lane 1 : K562 cell line; Lane 2 : UT7 cell line, lane 3 : UT7-11 cell line expressing BCR-ABL; Lane 4 and 5 : CML IPSC line PB32 with and without Imatinib; (lane 6 to 9) in control embryonic stem cell line H1 and non Ph+ control IPS cell line PB33 with and without imatinib. **(B)** Expression profile of BCR-ABL gene by qPCR in CML IPS PB32 with and without Imatinib.

### BCR-ABL signaling in CML iPSC

The next question we asked was to determine if at the pluripotent stage, BCR-ABL-expressing CML iPSC were able to generate a tyrosine kinase activity, by either autophosphorylation or by phosphorylation of the Crk-L protein. To evaluate this hypothesis, we used Western blots using the pTYR4G10 phosphotyrosine antibody. As can be seen in Figure 2A, PB32 cells showed a highly phosphorylated BCR-ABL p210 protein as compared to control IPSC. Furthermore, this phosphorylation level corresponding to p210 BCR-ABL was decreased after imatinib treatment suggesting the detection of an autophosphorylated BCR-ABL in PB32 cells (**figure 2A**). We then looked for the phosphorylation of the surrogate BCR-ABL targets CrkL. We used an antibody against its phosphorylated form in CML-IPSC (4 independent experiments). As can be seen in Figure 2B, Tyrosine phosphorylated CrkL proteins were present in CML-IPSC and the level of them was significantly decreased after imatinib treatment (Mann Whitney pvalue 0,03). We confirmed this observation with ImageJ quantification relative to the level of b-ACTIN (**figure 2B**). These results therefore suggested that BCR-ABL signaling is preserved in CML iPSC, at least as assessed by its phosphotyrosine generation potential.

**Figure 2:**
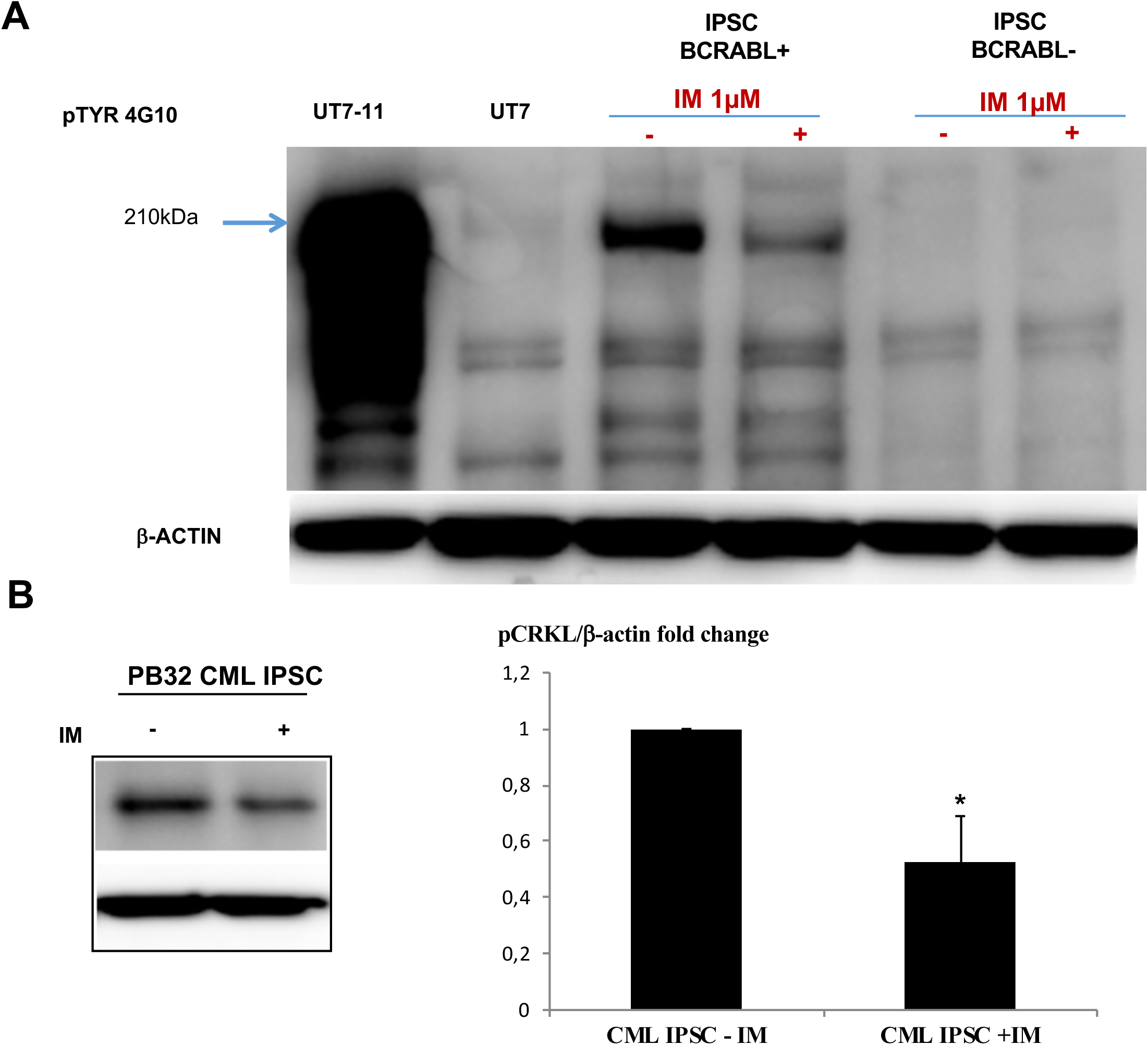
Evaluation of BCR-ABL phosphorylation in CML iPSC line PB32. **(A)** Western blot analysis of BCR-ABL phosphorylation levels with a phospho-tyrosine antibody (4G10 clone). Lane 1 and 2 : BCR-ABL-positive cell line UT7-11 and its negative counterpart control UT7; Lane 3 and 4 : PB32 CML iPSC with and without Imatinib; Lane 5 and 6: A control iPSC line without Ph1 chromosome, cultured with and without Imatinib. **(B)** ImageJ quantification of pCRKL protein in CML IPS cell line with and without Imatinib.

### CML-derived iPSC generate high numbers of blast CFC *in vitro*

Bl-CFC, generated according to the technique previously published (Lu et al., 2007) were quantified in the methylcellulose from day 5 to day 7. At day 5, we observed different types of small, either tight or diffuse colonies comprised of uniformly round cells corresponding top Bl-CFC (**figure 3B**). As compared to H1 control, we observed a 4-fold increase of the Bl-CFC clonogenic potential in the IPSC derived from CML (unpaired t.test with Welch’s correction p= 0.0037) in > 3 independent experiments in duplicate) (**figure 3A**). The control human IPS cell line PB33 failed to generate Bl-CFC in most experiments.

**Figure 3:**
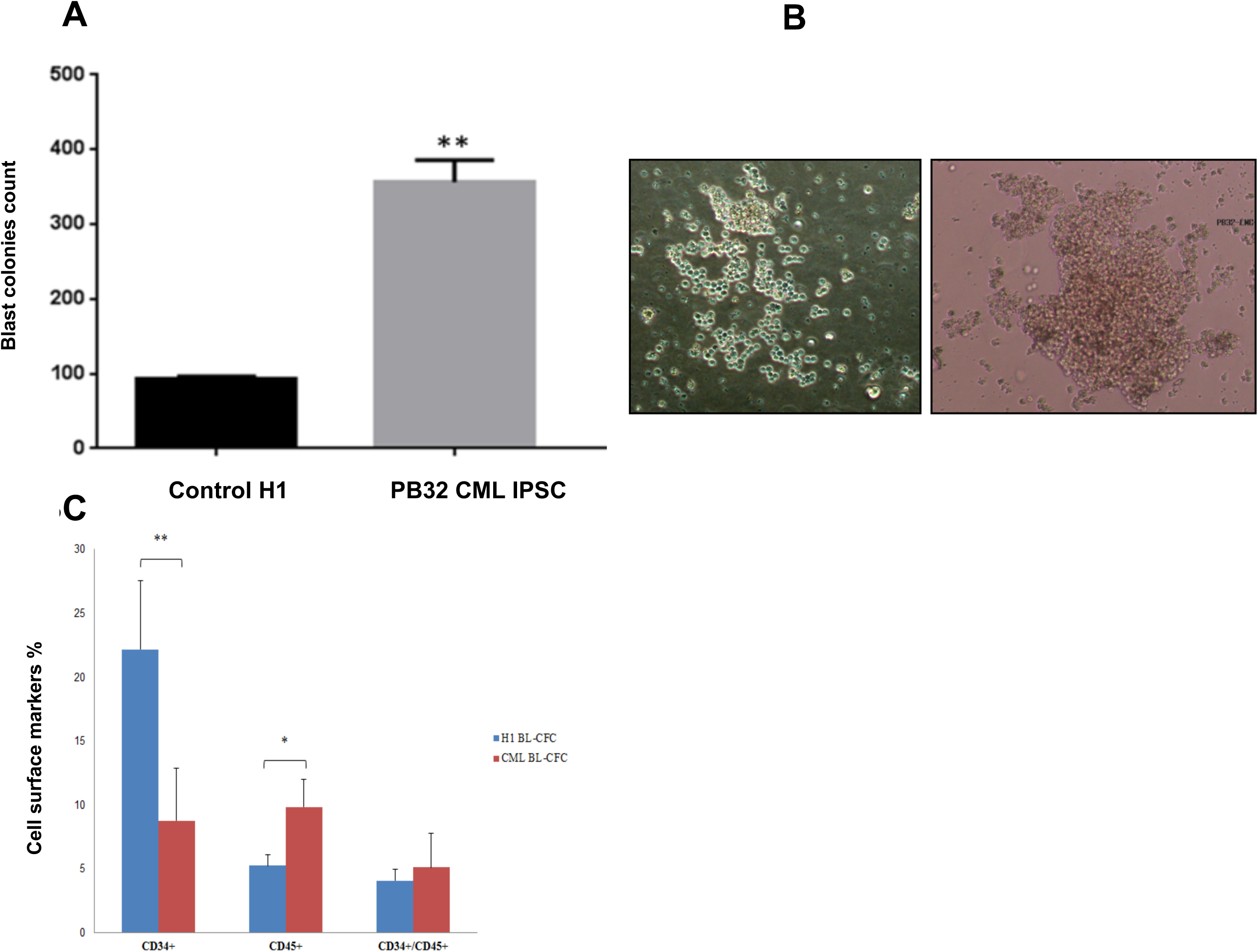
Impact of BCR-ABL expression in hematopoietic potential of CML IPS derived Bl-CFC. **(A)** Blast colony forming cells (Bl-CFC) derived from control embryonic stem cell line H1 and PB32 CML iPSC. **(B)** Representative photos pf Bl-CFC. **(C)** Flow cytometry analysis of hematopoietic markers CD34 and CD45 in Bl-CFC derived from control ESC (H1) and PB32 CML iPSC.

To characterize these blast cell colonies, we performed FACS analysis after collecting day 5 Bl-CFC from methylcellulose cultures (n= 4 experiments). As compared to Bl-CFC generated from H1-ESC, Bl-CFC derived from CML-IPSC expressed significantly less CD34 marker (mean % of 9% versus 22% in H1 p = 0.0069) and significantly more CD45 (10% vs 5%, respectively (p= 0.0156)(**figure 3C**). These results suggested that CML Bl-CFC had higher proliferative potential and were more committed to hematopoietic lineage.

In addition, FACS analyses of CML-Bl-CFC showed expression of both hematopoietic and endothelial markers (**supp figure 1**). These cells co-expressed CD309, VEGF-R, CD34, as well as CD31. Cells co-expressing the VE-cadherin (CD144) with CD309, BB9 or CD34 were also present. In addition, a clear hematopoietic nature of these cells was identified by the generation of CD45+ cells expressing CD43, CD34, were detected.

We also showed that CML Bl-CFC expresses IL1RAP which has been shown to be expressed in the primitive CML stem cells (Landberg et al., 2016). IL1-RAP+ cells co-expressed CD45.

### Bl-CFC and embryoid body-mediated hematopoietic clonogenic cell potential is increased in CML-iPSC

We performed hematopoietic differentiation using CFU assay from Bl-CFC or from EB (n= 3 experiments) (**figure 4**). These experiments showed a major and reproducible increase of CFC number EB and Bl-CFC generated from PB32 as compared to H1cell line, which is a reference for its hematopoietic potential (Kaufman et al., 2001). As a matter of fact, seeding of 10×10^3 CML Bl-CFC, generated approximately 5 times more hematopoietic colonies of all types (**figure 4A-B**) as compared to those produced from H1 Bl-CFC. Similarly, seeding of PB32 EB’s generated 5 to 8 times more hematopoietic colonies as compared to control EB, this increase being highly significant (ANOVA test with multiple comparison on PRISM software pvalue <0.0001)

**Figure 4:**
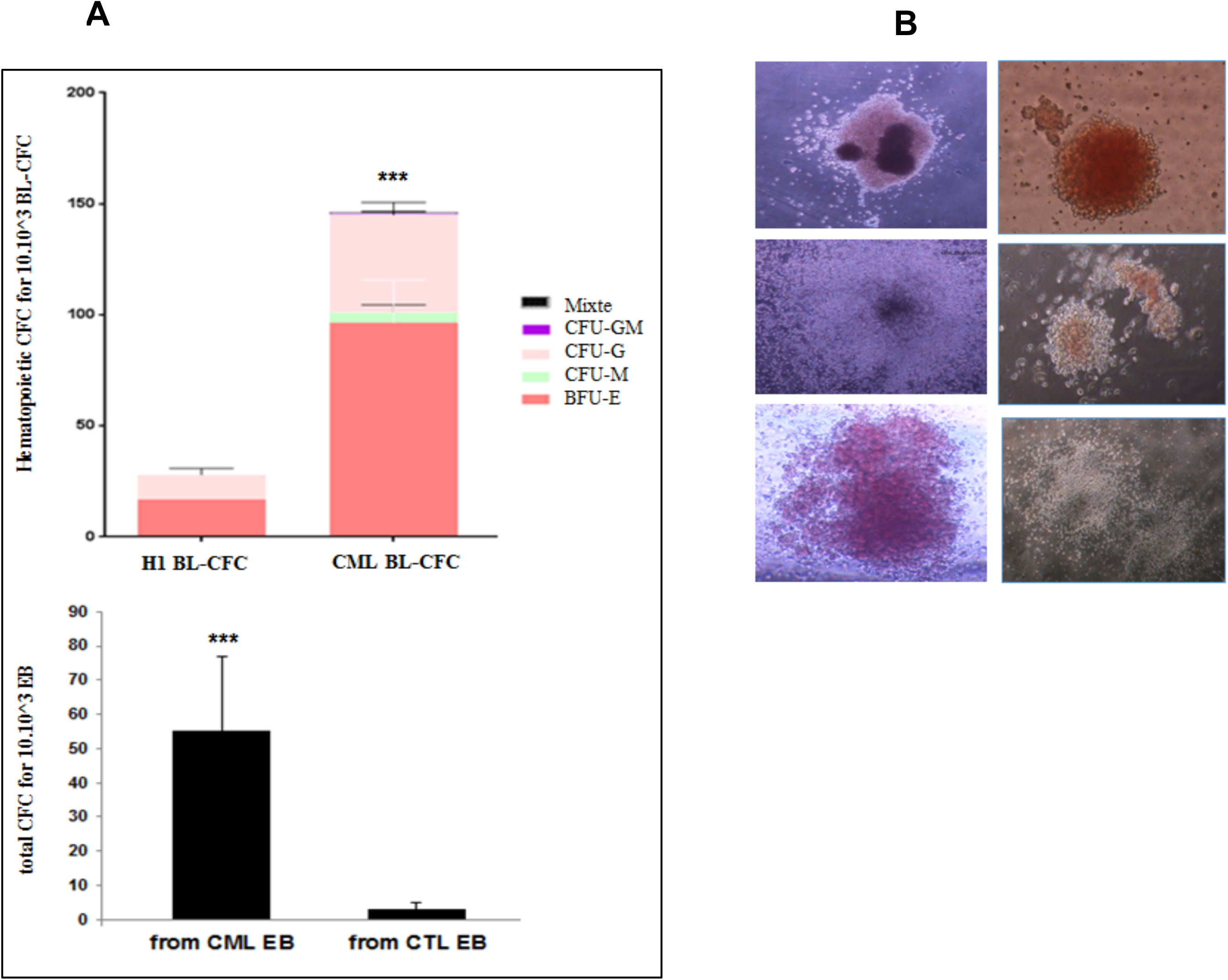
Hematopoietic differentiation from Bl-CFC and EB derived from PB32 CML iPSC. **(A)** Colony forming cell (CFC) count from 10.10^3^ control and CML Bl-CFC or EB. **(B)** Photos of hematopoietic cell colonies.

One interesting finding was the fact that we have observed constantly a higher hematopoietic potential when Bl-CFC were used as compared to EB-derived hematopoiesis (**Figure 4A**)

### Evaluation of BCR-ABL expression in progenitor cells

As we have generated Bl-CFC and EB-derived hematopoietic cells from the iPSC expressing BCR-ABL as described above, the next question was to determine if these primitive cells expressed also BCR-ABL after differentiation. These experiments showed the expression of BCR-ABL mRNA in both Bl-CFC and EB (**figure 5A**). This expression was confirmed in Western blots (**figure 5B**). CML-IPSC derived Bl-CFC expressed significantly more BCR-ABL mRNA than that observed in EB (unpaired t.test with Welch’s correction p = 0,0006) (figure 5A).

**Figure 5:**
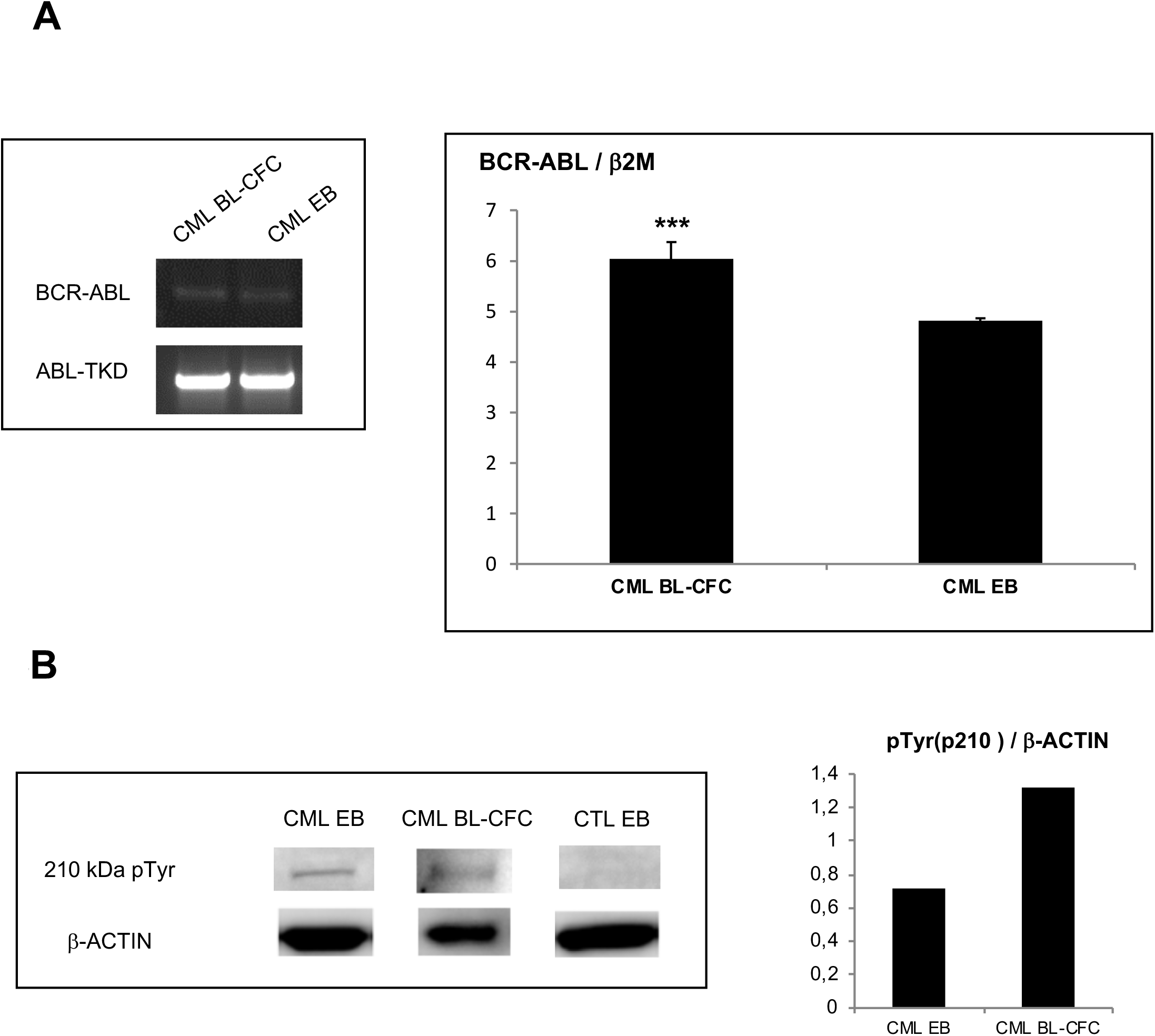
BCR-ABL expression in embryoid bodies (EBs) and Bl-CFC cells derived from CML iPSC. **5A (Left panel) :** PCR analysis to detect BCR-ABL in CML iPSC-derived blast colony forming cells (Bl-CFC) and embryoid bodies (EB). **5A (Right panel)** : Expression profile of BCR-ABL gene by RT-qPCR in CML iPSC-derived Bl-CFC and EB. **5B (Left panel):** Western blot analysis to detect BCR-ABL phosphorylation level using a phospho-tyrosine antibody. 5**B (Right panel):** ImageJ quantification of pTYR in iPSC-derived EB and Bl-CFC.

The use of P-Tyr antibodies showed also a higher expression of phosphorylated BCR-ABL, in CML Bl-CFC (**figure 5B**).

### Evaluation of the integrity of the BCR-ABL signaling at the level of embryoid bodies

The next question we have addressed was to determine if at the stage of EB’s where hematopoietic commitment occurs (Keller et al., 1993), the tyrosine kinase activity of BCR-ABL was sufficiently active to phosphorylate the surrogate marker Crk-L. To determine this event, we looked for the phosphorylated form of BCR-ABL (pBCR-ABL) using western blot analysis. As shown in Figure 6A the phosphorylated p210 band was detected only in our PB32-EB as well as our BCR-ABL-expressing cell line UT7-11 (**Figure 6A**). The use of Imatinib induced a major decrease of pBCR-ABL band in PB32 EB (**Figure 6A**, lane 6). Similarly, phospho-CrkL expression was highly increased in PB-32 EB’s protein extracts, (**Figure 6A**, lane 5) and CrkL phosphorylation was reduced upon IM treatment of the cells (**Figure 6A**, Lane 6 and **Figure 6B**). Interestingly, Imatinib also induced a decrease of CrkL phosphorylation in CTLs EB (**Figure 6A**, Lanes 8 and 10 and **Figure 6B**).

**Figure 6:**
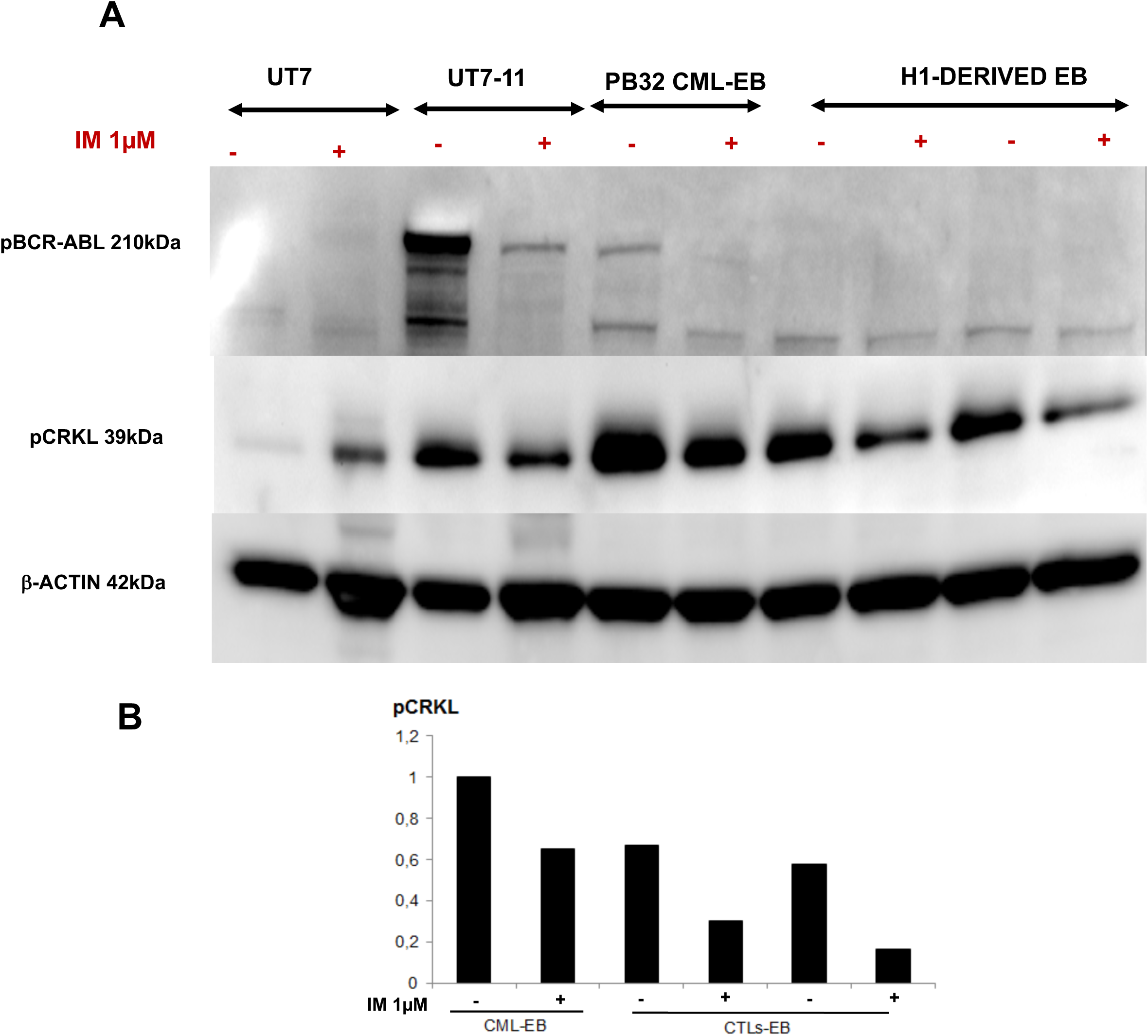
Western blot analyses of BCR-ABL phosphorylation levels and signalization after TKI treatment. **(A)** Western blot analysis of BCR-ABL phosphorylated protein and CRKL phosphorylation in CML and control EB with and without 1μM Imatinib compared to negative and positive control cell lines UT7 and UT7/11. **(B)** ImageJ quantification of pCRKL signal compared to β-Actin.

### Evaluation of the sensitivity of CML-iPSC-derived progenitors to tyrosine kinase inhibitors

As BCR-ABL was functional in our CML-iPSC at different stages of differentiation, we next asked whether we could determine if TKI could influence the growth of hematopoietic progenitors. We have then performed CFC assays in the presence or in the absence of TKI Imatinib and Dasatinib. As can be seen in Figure 7, CML-EB derived CFC were highly increased as compared to control (**Figure 7**). The use of IM and Dasatinib reduced the CFC growth by approximately 50 % (n = 3) and this inhibition was highly significant (ANOVA p = 0.002) (**Figure 7**).

**Figure 7:**
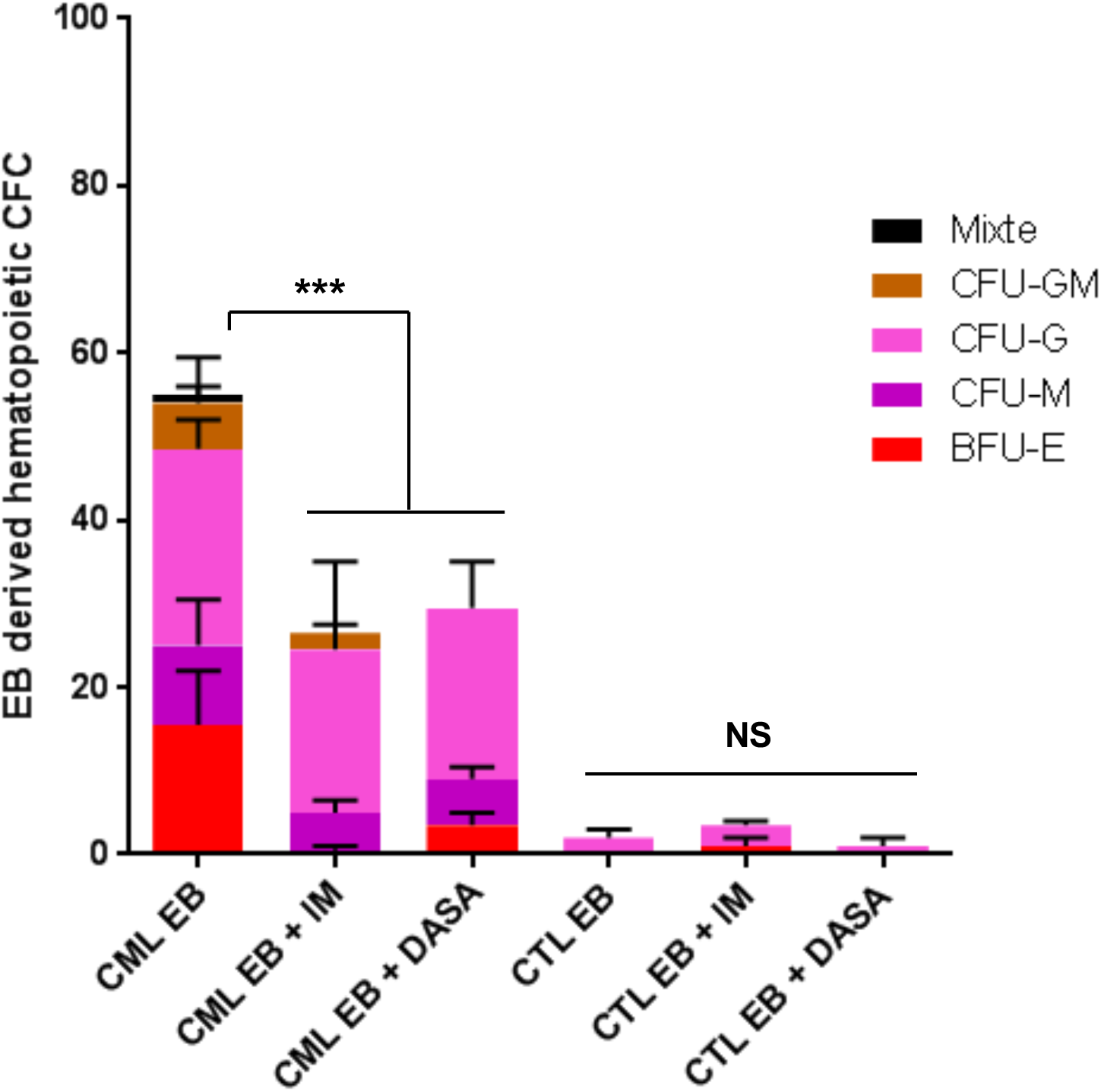
TKI sensitivity of PB32 CML iPSC and control hIPSC-derived hematopoietic cells. Hematopoietic cell colonies (CFC) count from CML and EB derived from control hIPSC (PB33) with and without 1μM Imatinib or 5nM Dasatinib.

### Further characterization of CML-iPSC derived hematopoietic cells by expression of genes involved in hematopoietic development

Hematopoietic commitment in adult is under the control of a well-defined set of genes. We analyzed the expression of some of these genes during differentiation from IPSC to Bl-CFC or EB derived hematopoietic progenitors (**figure 8 gene expression** and Table 1). As shown in Figure 8, the expression of PU.1, Gfi.1 and C-myb was found to be increased in Bl-CFC compared to EB. Furthermore, Ikaros and GATA3 which are usually expressed in T lymphocytes lineage are also highly expressed in H1-derived and CML derived Bl-CFC. Surprisingly, Lmo2 is only expressed in CML-IPSC derived Bl-CFC (**Figure 8**). CXCR4 expression as found to be downregulated in CML Bl-CFC or EB (**Figure 8**).

**Figure 8:**
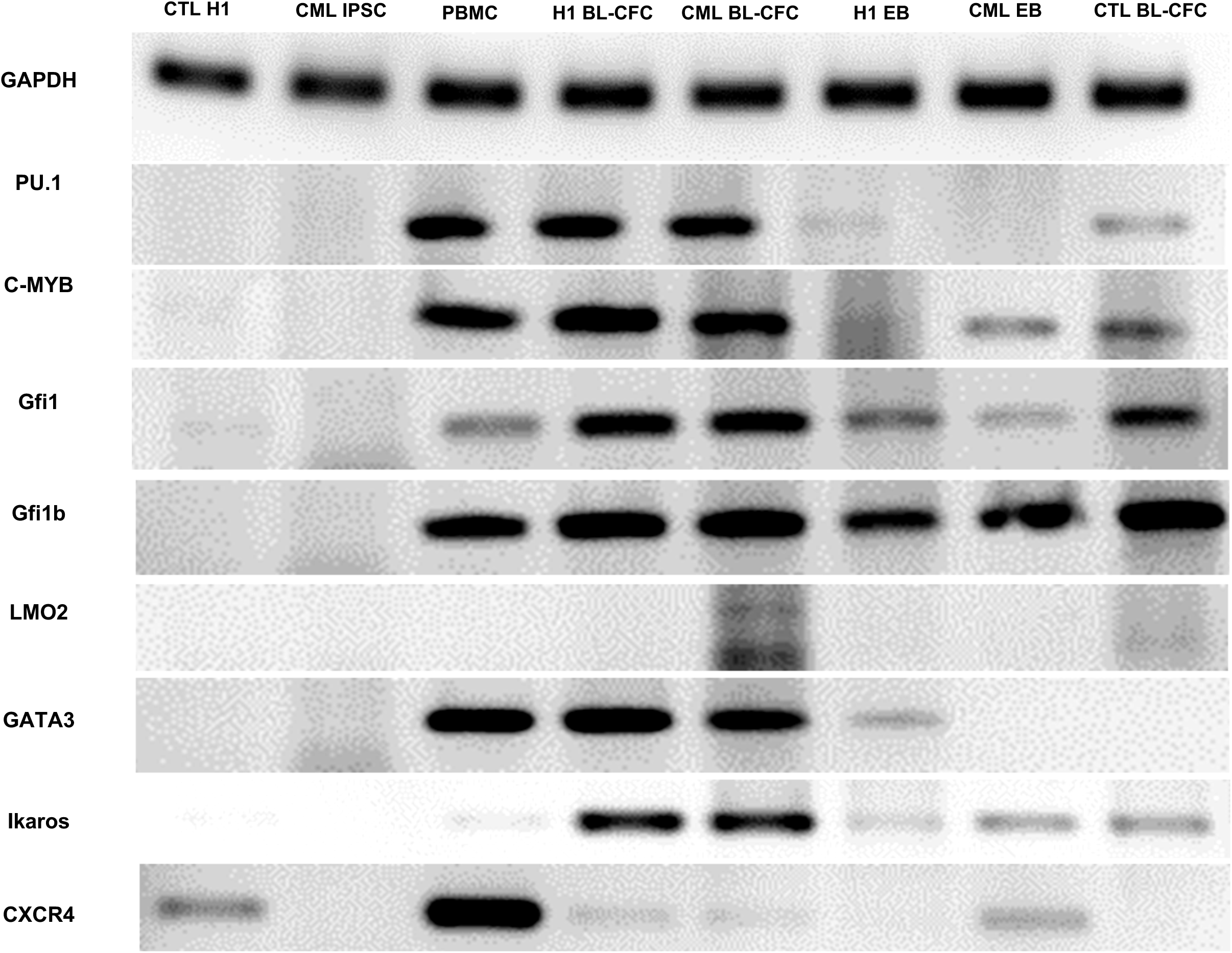
Hematopoietic gene expression profile. RT-PCR analysis of hematopoietic genes involved in hematopoietic development and differentiation. Lanes 1 : Human ESC cell line (H1) Lane 2 : PB32 CML iPSC; Lane 3 : Normal control PBMC; Lane 4 and 5 : BL-CFC derived from H1 cell line or PB32 CML IPSC; Lane 6 and 7: EB derived from H1 cell line and CML IPSC; Lane 8 : Bl-CFC derived from a control iPSC (PB33) derived from a normal donor.

**Table 1.** Expression of selected genes involved in hematopoietic development and differentiation.

|  | H1 IPS | PB32 IPS | PBMC | BL-CFC from H1 | BL-CFC from PB32 | EB from H1 | EB from PB32 | BL-CFC from PB33 |
| --- | --- | --- | --- | --- | --- | --- | --- | --- |
| LMO2 | - | - | - | - | ++ | - | - | +/- |
| GFI1 | - | - | +/- | ++ | ++ | + | +/- | ++ |
| GFI1b | - | - | ++ | ++ | ++ | + | + | ++ |
| PU.1 | - | - | ++ | ++ | ++ | +/- | - | + |
| GATA3 | - | - | ++ | ++ | ++ | +/- | - | - |
| C-myb | - | - | ++ | ++ | ++ | +/- | + | + |
| Ikaros | - | - | +/- | ++ | ++ | +/- | + | + |
| CXCR4 | + | - | ++ | +/- | +/- | - | + | - |
| BCR-ABL | - | + | - | - | + | - | + | - |

**Table 2:**
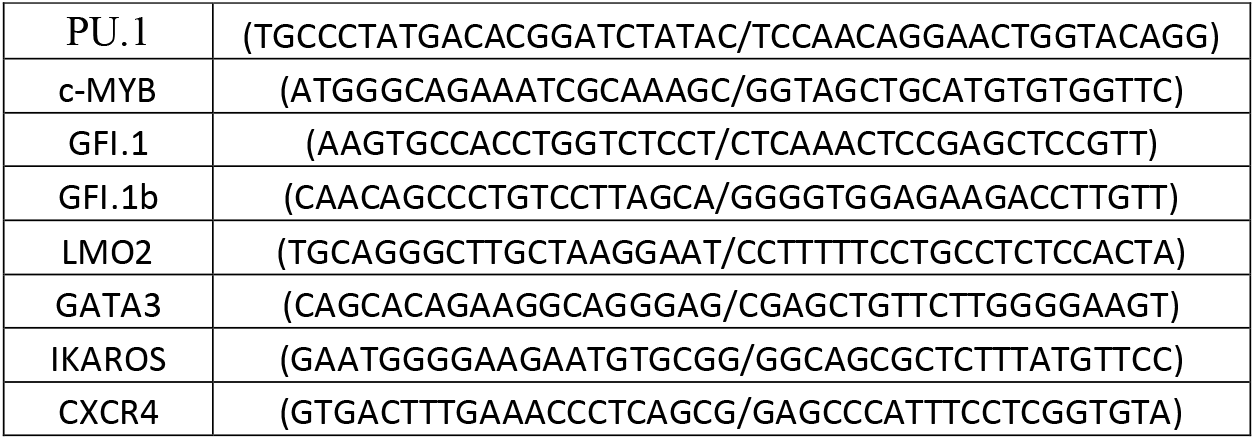
List of primers used for PCR analyses.

### Evaluation of AHR expression in CML and control pluripotent cells

Aryl hydrocarbon receptor signaling has been shown to be involved in hematopoietic cell cycle and particularly HSC quiescence (Singh et al., 2009). We have previously showed that AHR expression is downregulated in primary CML cells, potentially contributing to the myeloproliferative phenotype of CML (Gentil et al, 2018). We have therefore analyzed the expression of AHR in CML iPSC as compared to control iPSC. As can be seen in figure 9A, CML iPSC expressed significantly lower levels of AHR mRNA as compared to control iPSC but this level was equivalent to that observed in hES cell line H1 (n=3, p =0.0095) (**Figure 9A**).

**Figure 9:**
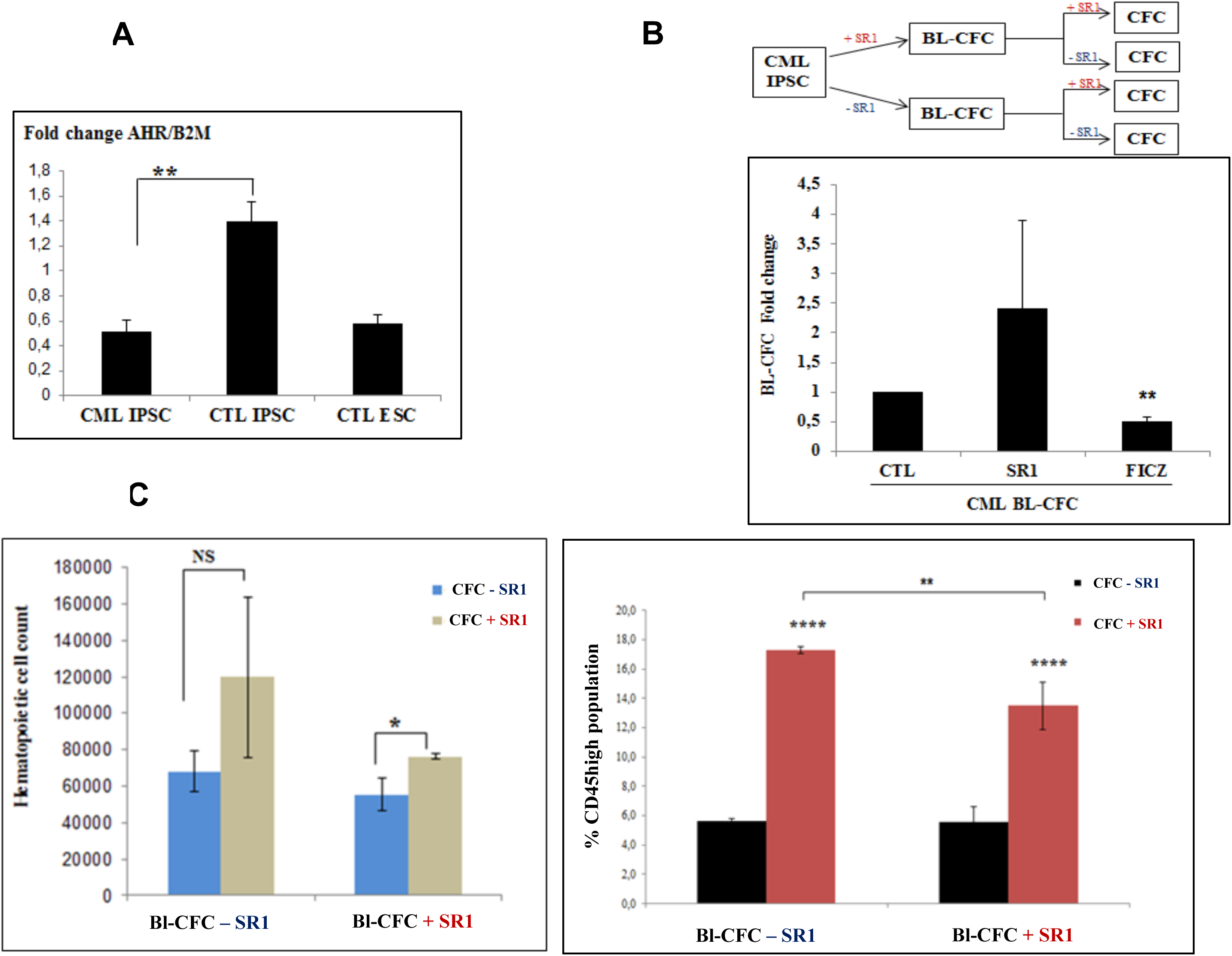
Modulation of hematopoietic potential by AHR signaling in CML iPSC-derived Bl-CFC. **(A)** AHR gene expression levels analyzed by RTqPCR in CML IPSC as compared to iPSC from normal donor (PB33, CTL IPSC) and human ESC line H1. **(B)** Schematic representation of the experimental approach. Upper panel : iPSC were used to generate Bl-CFC in the presence or in the absence of AHR antagonist SR1 for 5 days. Bl-CFCs generated in either way were then used to generate hematopoietic cells in the presence or in the absence of SR1. Lower panel: Evaluation of the effect of AHR antagonist SR1 or AHR agonist FICZ in the generation of CML Bl-CFC. **(C)** Impact of the AHR antagonist SR1 on the generation of CD45+ hematopoietic cells according to the presence or absence of SR1 during Bl-CFC cultures (see text).

### Evaluation of the effects of AHR antagonism

Inhibition of AHR signaling by the use of AHR antagonist StemRegenin (SR1) has been shown to lead to an expansion of normal hematopoietic progenitors (Boitano et al., 2010) as well as CML progenitors (Gentil et al, 2018). To determine the effects of AHR antagonism in the context of CML iPSC expressing high levels of AHR mRNA, we have performed experiments at two different stages of differentiation, as described in Figure 9B. As can be seen in this Figure, we generated Bl-CFC either in the presence or in the absence of SR1 added to the cultures (Fig 9B). From Bl-CFC generated this way, we have then performed hematopoietic differentiation using CFC assays in the presence or in the absence of SR1 added in methylcellulose cultures.

We first analyzed the effects of SR1 on the Bl-CFC potential of PB32 CML iPSC. As can be seen in Figure 9B, there was an increase of Bl-CFC numbers when SR1 was added to the cultures during Bl-CFC generation (Fig 9B). We next analyzed the potential of the Bl-CFC generated using SR1 towards hematopoietic differentiation. Quantitative and qualitative analyses of hematopoietic cells generated in these cultures in the absence or in the presence of SR1 is shown in Figure 9C. For these experiments cells were collected from methylcellulose cultures at day +14, counted and washed before staining with a CD45 antibody.

As can be seen in Figure 9C, there was a significant increase of CD45+ hematopoietic cell proliferation only when SR1 was added in CFC cultures without requirement of SR1 addition during Bl-CFC generation (n= 3 Experiments) (**Figure 9C**). After phenotypic characterization, we also showed that SR1 addition yielded increased numbers of hematopoietic cells expressing CD45+CD33+ (Suppl Fig 2). Cells with CD45+CD31+ markers were also increased by AHR signaling inhibition (Suppl Fig 2).

## DISCUSSION

The elusive nature of the most primitive CML stem cells which are very difficult to study in human samples stimulated a major research during the last 10 years to generate models of CML stem cells (Sloma et al., 2010). As matter of fact, primary CML stem cells which are very difficult to isolate and expand, are at the origin of relapses upon discontinuation of TKI therapies (Lee et al., 2016; Mahon et al., 2010; Ross et al., 2013; Chomel et al, 2011; Chomel et al, 2016). Several murine CML models have been developed to this purpose but they do not recapitulate faithfully the most primitive human CML cells. The use of embryonic stem cell models are of interest as the overexpression of BCR-ABL has been shown to be a major stimulus for generation of long-term leukemic hematopoiesis from murine ES cells (Melkus et al., 2013; Perlingeiro et al., 2001; Peters et al., 2001). However, one of the limitations of the ES models is the need to overexpress BCR-ABL under the control of different promoters potentially leading to results not relevant to the pathophysiology of human CML. With the advent of iPSC technology, it became of major interest to use human CML cells for reprogramming with the goal of reproducing the earliest stages of human CML and potentially to use them for drug screening purposes. The first work published in this field showed the feasibility of reprogramming the human CML cell line KBM to pluripotency (Carette et al., 2010) with the possibility of generating BCR-ABL-expressing CD45+ cells. Other groups used primary CD34+ leukemic cells which have been shown to be induced to pluripotency using different methodologies (Bedel et al., 2013; Kumano et al., 2012). The usefulness of this technology lies on the fact it becomes possible to obtain an unlimited numbers of primitive cells using the background of the primary leukemic cells in any given patient. However, experiments aiming to generate increased hematopoietic potential have been disappointing. In fact some groups showed that, CML iPSC expressed very low levels of BCR-ABL and did not generate a myeloproliferative phenotype (Bedel et al., 2013; Kumano et al., 2012). One group showed the possibility of using CML iPSC to discover a novel signaling pathway operational in human CML (Suknuntha et al., 2015).

The iPS cell line used in the experiments reported here was generated from the primary leukemic cells of a 14-year old patient with TKI-resistant leukemia (Telliam et al 2016). This patient had no evidence of ABL-kinase domain mutation and the reason of Imatinib résistance was not determined. The first goal of the reprogramming of the leukemic cells of the patient was to determine if BCR-ABL expression and induced signaling could be intact at different stages. We show here that BCR-ABL expression is present at the most primitive iPSC stage and BCR-ABL autophosphorylation is present in this context as it would have been expected in an adult leukemic cell context. Similarly, we show here that during the differentiation process through the stages of hemangioblasts and EB’s, BCR-ABL expression remains intact with evidence of tyrosine-phosphorylation of both CML Bl-CFC and EB’s (Figure 5B). In CML EB’s, the downstream target Crk-L is phosphorylated (Figure 6A, 6B) and BCR-ABL phosphorylation is reduced by IM treatment of cells (Figure 6A). This allowed us to perform TKI sensitivity experiments showing that these cells indeed respond to TKI inhibition at the level of CFC and this inhibition was seen with both IM and Dasatinib (Figure 7). The patient at the origin of the iPSC was shown to be clinically resistant to Imatinib but the sensitivity of his cells in CFC experiments was not performed at diagnosis.

To further characterize these iPSC, we have analyzed the expression of hematopoietic genes involved in hematopoietic development at different stages of differentiation as compared to H1 ESC and adult hematopoietic cells. As can be seen in Figure 8, CML iPSC at the pluripotent stage did not express any of the hematopoietic genes involved in hematopoietic development (PU1, c-MYB, Gfi1, Gfi1b, LMO2, GATA3, Ikaros) nor CXCR4 (Figure 8). The expression of these genes appeared during differentiation and as compared to H1-derived EB’s, CML EB’s showed a slight increase of Gfi1b, CXCR4 and c-MYB expression (Figure 8). At Bl-CFC stage, the expression of GATA3 was found to be higher as compared to EB stage and interestingly LMO2 expression was detectable only in CML Bl-CFC (Figure 8). Interestingly, c-MYB has been shown to play a role in BCR-ABL-induced leukemogenesis (Lidonnici et al., 2008).

Our data show here also for the first time to our knowledge, that AHR pathway which we have shown to be downregulated in adult CML cells is also downregulated in CML iPSC. It is not known at this time if this downregulation is specific to CML-iPSC, as ES cells have been shown to exhibit a downregulation of AHR under the influence of OCT4 (Kang and Wang, 2015). We have used the antagonism of this pathway to show that it is possible to further expand hematopoietic cells of the CML iPSC by this strategy. As we have shown in a previous work, data reported here suggest also that AHR pathways could represent a druggable target as the AHR agonist FICZ inhibited the Bl-CFC growth (Figure 9B). This inhibitory effect was previously reported using primary CML stem cells (Gentil et al, 2018).

In summary, we have generated and characterized a unique CML iPSC from of TKI-resistant patient’s leukemic cells. These cells partly recapitulate some features of the disease with essentially a myeloproliferative phenotype. This phenotype could be modulated by the antagonism of the AHR pathway which we have shown to be efficient in adult primary CML cells. IPSC-derived cells express BCR-ABL able to tyrosine-phosphorylate downstream targets and this phosphorylation is responsive to TKI inhibition. Finally further experiments will test the possibility of generating long-term hematopoiesis from these patient-specific iPSC after transplantation of iPSC-derived EB’s in immunodeficient mice.

## SUPPLEMENTAL DATA

**Supplemental Figure 1:**
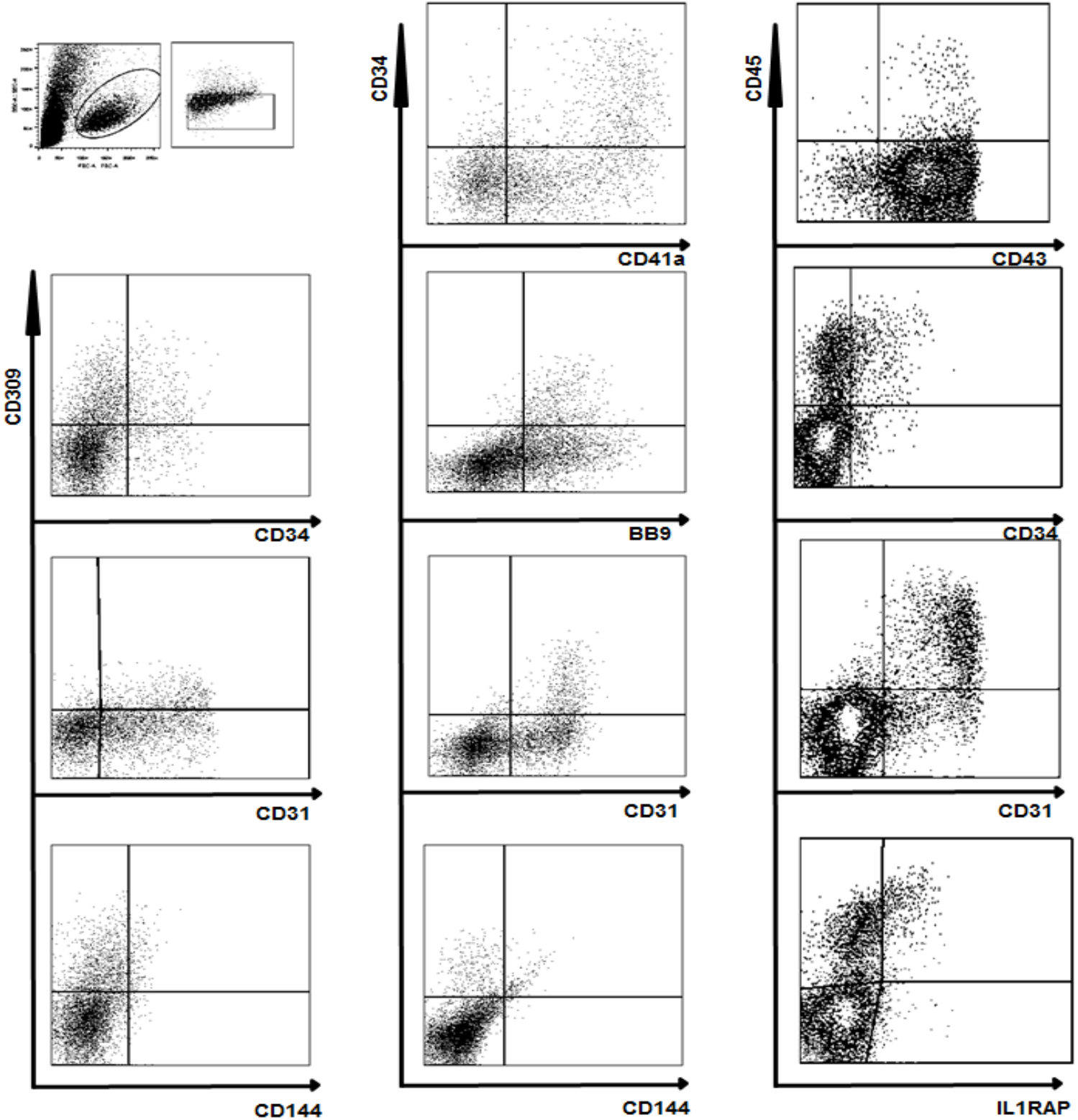
Phenotypic characterization of Bl-CFC derived from CML-iPSC, with evidence of endothelial differentiation (CD31, CD309).

**Supplemental Figure 2:**
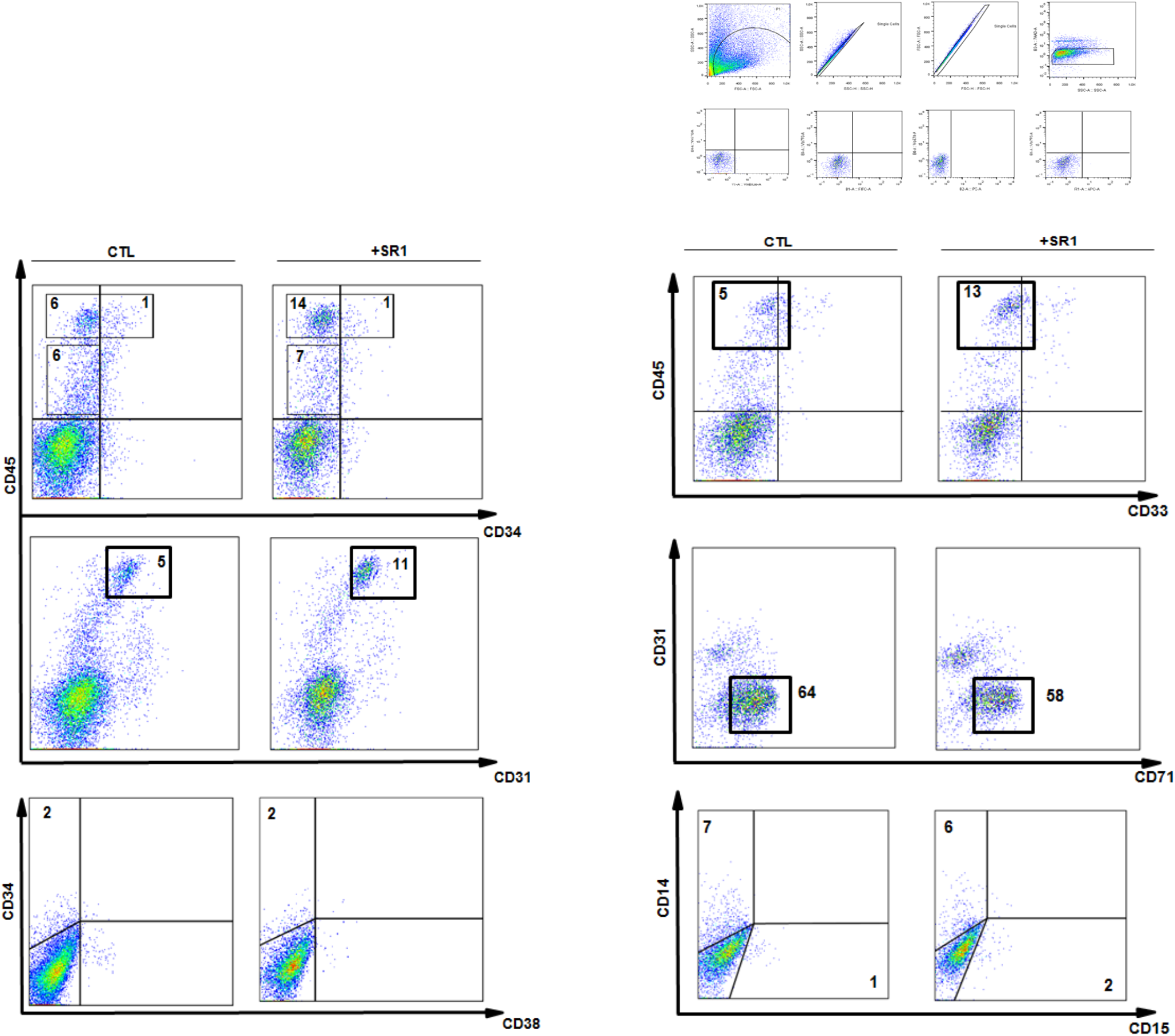
Phenotypic characterization of PB32-CML iPSC-derived hematopoietic CFC, cultured in the presence or in the absence of SR1 for 14 days (see text).

